# Pf Bacteriophage Inhibits Neutrophil Migration in the Lung

**DOI:** 10.1101/2022.10.12.511980

**Authors:** Medeea C. Popescu, Nina Pennetzdorfer, Aviv Hargil, Gernot Kaber, Paul L. Bollyky

**Affiliations:** Division of Infectious Diseases and Geographic Medicine, Stanford University, Stanford, CA 94305, USA; Immunology Program, Stanford University

## Abstract

Bacteriophages are abundant in the human body, including at sites of infection. We report that Pf4 phage, a filamentous bacteriophage produced by *Pseudomonas aeruginosa*, dampens inflammatory responses in response to either *P. aeruginosa* airway infection in a mouse model of acute pneumonia or bacterial endotoxin treatment. Pf4 triggers TLR3-dependent type I interferon production and antagonize production of anti-bacterial cytokines and chemokines. In particular, Pf4 phages inhibit CXCL5, preventing efficient neutrophil chemotaxis in response to endotoxin. These results suggest that Pf4 phages alter innate immunity to bacteria potentially dampening inflammation and neutrophil migration at sites of bacterial colonization or infection.

## INTRODUCTION

Bacteriophages, viruses that parasitize bacteria, are abundant at sites of bacterial colonization and infection, including in the lungs of patients infected with *Pseudomonas aeruginosa*. However, it remains unclear how the innate immune system senses phages or how phages impact sensing of bacteria (*Popescu* et al., 2021). While some phages are potent immunogens and have been used for vaccine development (*Dąbrowska* et al., 2014; *de Vries* et al., 2021), other phages induce minimal inflammation, including phages used in phage therapy (*Cano* et al., 2021; *Liu* et al., 2021). The mechanisms that drive these distinctions are unclear.

We and others have previously investigated the impact of Pf phages on their bacterial and human hosts. Pf is a genus of filamentous bacteriophages that infect the common bacterial pathogen *P. aeruginosa* and includes the phages Pf1-Pf8 (*Knezevic* et al., 2015; *Mai-Prochnow* et al., 2015; *Hay* and *Lithgow*, 2019; *Fiedoruk* et al., 2020). Pf phages are Inoviruses and have a single stranded (ss) DNA genome packaged within a helical filamentous structure made up of thousands of copies of the major coat protein CoaB (*Marvin* et al., 2014; *Hay* and *Lithgow*, 2019; *Roux* et al., 2019). Pf virions are ~ 6-7 nm in diameter and ~ 1-2 μm in length (*Janmey* et al., 2014). Approximately half of *P. aeruginosa* isolates harbour Pf phages (*Knezevic* et al., 2015; *Burgener* et al., 2019). Unlike lytic or lysogenic phages that lyse their bacterial hosts after replication, Pf phages follow a chronic infection life cycle whereby Pf virions are continuously extruded from the bacterial cell envelope without lysis (*Secor* et al., 2020).

Production of Pf4, a well-studied member of the Pf family, contributes to bacterial phenotypes associated with *P. aeruginosa* chronic infection, including reduced twitching motility, increased adhesion, enhanced biofilm formation, and antibiotic tolerance (*Rice* et al., 2009; *Secor* et al., 2015, 2017; *Sweere* et al., 2019; *Tarafder* et al., 2020). Indeed, filamentous phages like Pf4 contribute to the fitness of their bacterial hosts and the pathogenesis of *P. aeruginosa* infections (*Mai-Prochnow* et al., 2015; *Bille* et al., 2017; *Pant* et al., 2020; *Pourtois* et al., 2021; *Schmidt* et al., 2022). Consistent with this, we previously reported that Pf phages are common in individuals with cystic fibrosis (CF) and that *P. aeruginosa* infections associated with Pf+ strains are characterized by older age, advanced lung disease, worse disease exacerbations, and antibiotic resistance in this group (*Burgener* et al., 2019). Pf phages are likewise associated with chronic *P. aeruginosa* wound infections (*Sweere* et al., 2019).

Along with effects on bacterial pathogenesis, Pf4 phage enables *P. aeruginosa* to evade the mammalian host immune response. In a model of acute lung infection, mice inoculated with *P. aeruginosa* supplemented with Pf4 phage did not develop sepsis, had less pulmonary inflammation, and survived significantly longer than mice infected with WT *P. aeruginosa* only (*Secor* et al., 2017). In a chronic wound infection model, Pf4 phage was associated with impaired phagocytosis and reduced levels of tumour necrosis factor alpha (TNF-α) (*Sweere* et al., 2019). These effects were mediated by Toll-like receptor 3 (TLR3) and type I interferon (IFN) production (*Sweere* et al., 2019). However, the nature of these responses was unclear given that Pf4 phage is a ssDNA virus and TLR3 recognizes double-stranded (ds) RNAs.

Here, we identify a role for Pf4 phage in inhibiting neutrophil chemotaxis in response to *P. aeruginosa* infection. We find that Pf4 phage downregulates the expression of neutrophil chemoattractants by human macrophages, resulting in impaired ability to recruit cells to the site of infection.

## MATERIAL AND METHODS

### Bacterial strains and growth conditions

Bacterial strains and plasmids used in this study are listed in the “Key Resources Table”. The *P. aeruginosa* isolate mPAO1 served as a wildtype that harbours the filamentous phage Pf4 in all experiments (*Rice* et al., 2009). Isogenic phage-free strain PAO1ΔPf4 was derived from strain PAO1 that lacks Pf4 entirely but can be reinfected by Pf4 (*Rice* et al., 2009). Unless stated otherwise, all *P. aeruginosa* strains were grown with aeration in Brain Heart Infusion (BHI, BD Bacto™, Cat. No. 237200), or Luria-Bertani (LB Miller Broth, BD Bacto™, Cat. No. DF0446-07-5) with aeration at 37°C. *Escherichia coli* DH5α was used for plasmid maintenance and was grown in LB with aeration at 37°C. Antibiotics were used in the following final concentrations: gentamycin (10 μg/ml for plasmid maintenance in *E. coli;* 100 μg/ml for plasmid maintenance in *P. aeruginosa*, VWR, Cat. No. 97062-974).

### Preparation and quantification of bacteriophages

*P. aeruginosa* mPAO1 was used to produce Pf4. Mock phage preparations from PAO1ΔPf4 were prepared according to the same protocol as other phage samples. Cultures were inoculated with an OD_600_ of 0.1 and grown to the mid-log phase in which the respective phage was added and coincubated overnight. The next day, the bacteria were harvested at 6,000 x *g*; 30 minutes; 4°C. The supernatant was sterile filtered using a 0.22 μm bottle top filter and treated with 50 U/ml of benzonase nuclease (Sigma Aldrich, Cat. No. E1014-25KU) for 2 hours. The phage solution was incubated with polyethylene glycol 8000 (PEG-8000, Sigma-Aldrich, Cat. No. P2139-500G) at 4°C overnight as described previously (*Boulanger* et al., 2009) and dialyzed against PBS for 48 hours. Purified phage preparations were quantified by plaque assays as well as by qRT-PCR using primers directed against the major coat protein of the respective phage and a vector based (pBS-SK back bone) standard comprising the gene encoding for the respective major coat protein. Phages were stored in 1x PBS at 4°C. Endotoxin was quantified by EndoZyme II assay (BioVendor, Cat. No. 890030) as per manufacturer’s directions.

### Tissue culture

Human U937 macrophages (ATCC, Cat. No. CRL-1593.2™) were maintained in RPMI (Thermo Fisher Scientific, Cat. No. 11875093) supplemented with 10% FBS (RMBIO, Cat. No. FGR-BBT), penicillin-streptomycin (Fisher Scientific, Cat. No. MT30002CI) and sodium pyruvate (Thermo Fisher Scientific, Cat. No. 11360070).

### Cell stimulation

For cytokine release and reporter assays, cells were seeded at a density of 5×10^4^ cells/well in a 96-well plate. Unless otherwise specified, stimuli were as follows: 1×10^8^ pfu/well of purified phage preparation and 100 ng/ml LPS (Invivogen, Cat. No. tlrl-eblps). The TLR3 signalling inhibitor (EMD Millipore, Cat. No. 614310) was used at 27 μM and anti-human IFNAR2 antibody (PBL Assay Science, Cat. No. 21385-1) was used at 100 ng/ml. Cells were incubated at 37°C and 5% CO2 for the indicated timepoints, then centrifuged at 300 x *g* for 5 minutes prior to supernatant removal.

### Enzyme-linked immunosorbent assay (ELISA)

Human CXCL5 (BioLegend; Cat. No. 440904), human TNF-α (BioLegend; Cat. No. 430904) and human IFNβ (Invitrogen; Cat. No. 414101) were performed using manufacturer’s instructions. Absorbance was read on a Spark microplate reader (Tecan).

### Luminex analysis

U937 macrophages were plated in 96-well plates at 5×10^4^ cells/well. Cells were stimulated for 24 hours with 1×10^8^ pfu/well of purified Pf4 phage or equivalent volume ΔPf4 preparation and 100 ng/ml LPS (Invivogen, Cat. No. tlrl-eblps). Cells were centrifuged at 300 x *g* for 5 minutes and supernatant was collected and used for cytokine profiling through Luminex (The Human Immune Monitoring Center, Stanford). Human 89-plex kits were purchased from eBiosciences/Affymetrix and used according to the manufacturer’s recommendations with modifications as described below. Briefly: Beads were added to a 96 well plate and washed in a Biotek ELx405 washer. Samples were added to the plate containing the mixed antibody-linked beads and incubated at room temperature for 1 hour followed by overnight incubation at 4°C with shaking. Cold and room temperature incubation steps were performed on an orbital shaker at 500-600 rpm. Following the overnight incubation plates were washed in a Biotek ELx405 washer and then biotinylated detection antibody added for 75 minutes at room temperature with shaking. Plate was washed as above and streptavidin-PE was added. After incubation for 30 minutes at room temperature wash was performed as above and reading buffer was added to the wells. Each sample was measured in duplicates. Plates were read using a Luminex 200 instrument with a lower bound of 50 beads per sample per cytokine. Custom assay Control beads by Radix Biosolutions were added to all wells.

### Luminex data analysis

Technical replicate MFI values were averaged for each sample and transformed into log_2_ fold-change with respect to the control LPS sample. Data was plotted as a heatmap using the pheatmap package (v1.0.12) in R.

### Isolation and preparation of neutrophils

Human neutrophil isolation protocol was adapted from *Swamydas* et al. (2015). Briefly, whole blood was obtained from the Stanford Blood Center, diluted 1:1 in 1x PBS (Corning, Cat. No. 21-040-CV), and layered over a double Histopaque-based density gradient (density 1.119 g/ml (Sigma-Aldrich, Cat. No. 11191-6X100ML) and density 1.077 g/ml (Sigma-Aldrich, Cat. No. 10771-6X100ML). The suspension was centrifuged at 900 *x g* for 30 minutes at 22°C with no brake, and neutrophils from the interface of the Histopaque-1119 and Histopaque-1077 layers were collected and washed in RPMI + 10% FBS. Count and viability of neutrophils was determined by trypan blue exclusion.

### Neutrophil migration assay

Primary human neutrophils isolated as described were stained at room temperature for 20 minutes with 5 μM Calcein AM (BioLegend, Cat. No. 425201) and applied to the top well of a fluorescence-blocking 24-well transwell plate with 3 μm pores (Corning, Cat. No. 351156). Bottom wells were filled with conditioned media from U937 macrophages, and chemotaxis was tracked by plate bottom fluorescence using a Spark microplate reader (Tecan) over the course of 30 minutes.

### Murine pneumonia model

Experiments were performed as described in *Secor* et al. (2017). Briefly, an isolated colony of *P. aeruginosa* mPAO1 or PAO1ΔPf4 was grown to mid-exponential phase (OD600, 0.5) in 4 ml LB broth at 37°C with aeration. The mPAO1 culture was infected with 100 μl of Pf4 stock (10^10^ pfu/ml) to promote phage production. These cultures were grown overnight at 37°C with aeration. On the following day, bacteria were pelleted by centrifugation at 6,000 x *g* for 10 minutes, washed 3x in sterile PBS, and resuspended in PBS to a final concentration of 3×10^8^ CFU/ml. Mice (8-10wk C57BL/6, male; The Jackson Laboratory) were intranasally inoculated with 1.5×10^7^ CFU/ml in 50 μl PBS of each strain or PBS control.

Mice were sacrificed at 24 hours post infection. The lungs were lavaged with 0.8 ml sterile PBS as described previously (*Secor* et al., 2017). BAL fluid samples were pelleted, and supernatant was frozen for use in ELISA and qPCR assays. BAL cells were resuspended in 1 ml ACK lysis buffer 30 sec, followed by washing in PBS with 2% foetal calf serum and 1 mM EDTA (fluorescence-activated cell sorting [FACS] buffer). The cells were resuspended in 1 ml FACS buffer and stained with Zombie Near-IR Live/Dead (Biolegend, Cat. No. 423105), CD11b-Pacific Blue (Biolegend, Cat. No. 101223), CD11c-BV605 (Biolegend, Cat. No. 117333, CD45.2-BV650 (Biolegend, Cat. No. 109835), Ly6G-BV785 (Biolegend, Cat. No. 127645), Siglec F-PE (Biolegend, Cat. No. 155505), and Ly6C-AF647 (Biolegend, Cat. No. 128009) as per manufacturer’s instructions. After staining, the cells were fixed with 4% paraformaldehyde and then run on a Cytek Northern Lights machine, followed by analysis with FlowJo™ software. Total live cells within the CD45^+^ gate was reported, as well as live neutrophils within the CD45^+^ Ly6G^+^ gate. Absolute cell counts were adjusted by total BAL volume recovered.

### Statistical analysis

Where n is not stated, graphs show a representative experiment of n ≥ 3 assays, with n ≥ 3 technical or biological replicates. All statistical analyses were performed using GraphPad Prism (GraphPad Software, Inc. La Jolla, CA). Significance was assessed by ANOVA followed by Holm-Šídák test for multiple comparisons unless otherwise indicated. Depicted are means with standard deviation (SD) of the replicates unless otherwise stated. Statistical significance was considered *p* < 0.05. Non-significance was indicated by the letters “n.s.”. For experiments involving cell assays, each replicate was normalized against a specified control condition to control for varying interexperiment intensities of cytokine production or reporter protein secretion.

## RESULTS

### Pf4-overproducing *P. aeruginosa* elicits diminished neutrophil migration

Building on our previous finding that treatment with *P. aeruginosa* strain mPAO1 supplemented with Pf phage reduced pulmonary inflammation in response to bacterial infection (*Secor* et al., 2017), we examined *P. aeruginosa* infection in the absence of Pf4 phage. To this end, we treated mice with mPAO1 or an isogenic strain lacking Pf4, PAO1ΔPf4. These strains have comparable growth curves (*Sweere* et al., 2019) such that differences between them cannot be attributed to altered growth. After 24 hours, bronchoalveolar lavage (BAL) fluid was collected and evaluated for cell influx and chemokine/cytokine content (Figure 1A). Mice infected with PAO1ΔPf4 showed significantly greater neutrophil influx and pro-inflammatory cytokine production than mice infected with Pf4-producing mPAO1 (Figure 1B-C). mPAO1-infected mice also exhibited significantly lower levels of CXCL1, TNF-α, and CXCL5 in BAL as compared to PAO1ΔPf4-infected mice (Figure 1D-F).

**Figure 1:**
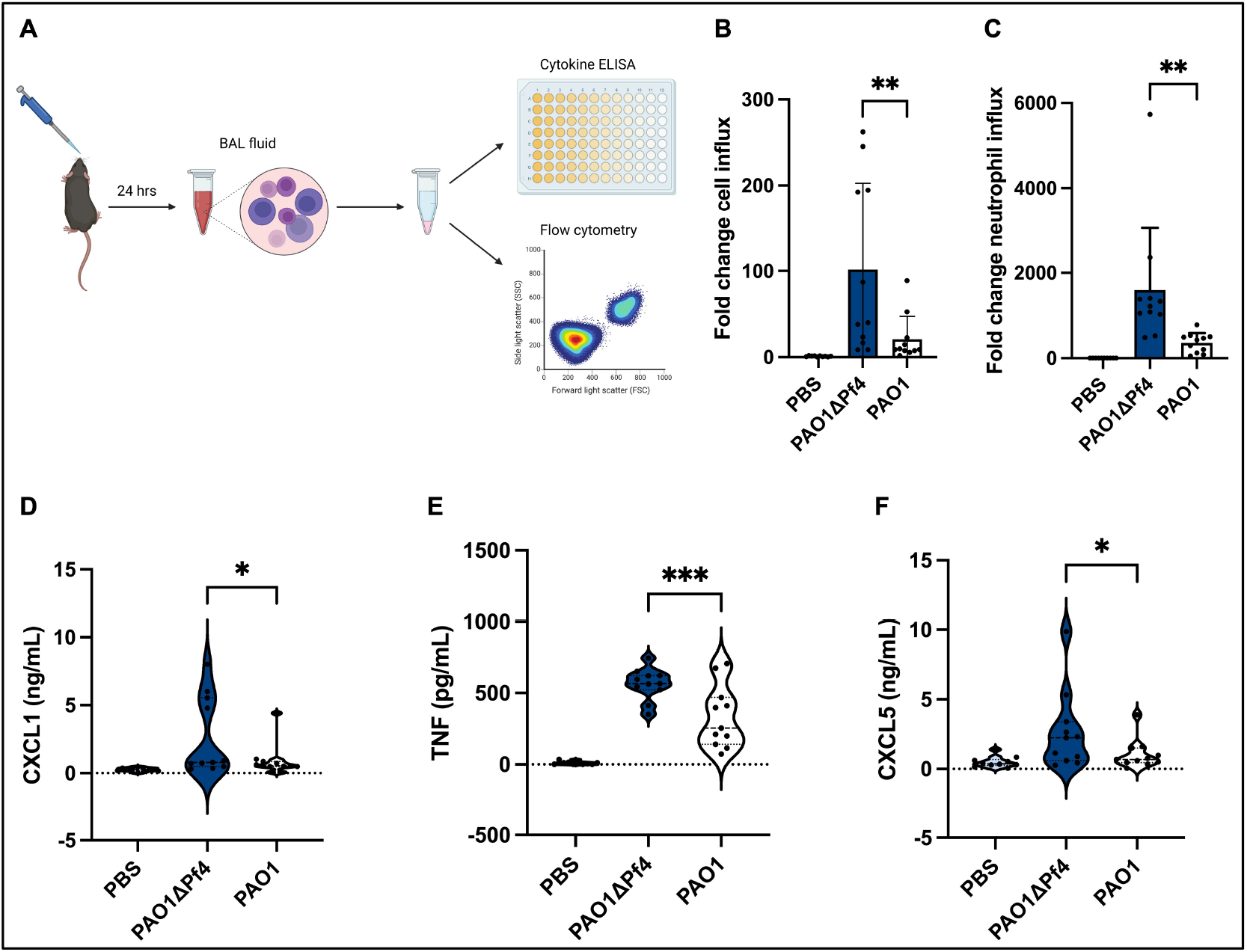
Neutrophil migration is reduced by Pf4 phage production in an acute *P. aeruginosa* pneumonia murine model. (A) Mice were inoculated intranasally for 24 hours with either PBS, mPAO1, or the isogenic *P. aeruginosa* strain PAO1ΔPf4 that lacks a functional copy of Pf4 phage. After 24 hours the animals were sacrificed and bronchoalveolar lavage (BAL) fluid was collected for cytokine measurements and cellular assessments via flow cytometry. (B and C) BAL fluids samples were evaluated for (B) total cell content and (C) neutrophil quantities as determined by flow cytometry. (D to E) BAL chemokine quantifications for (D) CXCL1, (E) TNF-α, and (F) CXCL5 as measured by ELISA. Data are displayed as the mean ± SD from three individual experiments with n ≥ 9 mice per experiment. * = *p* < 0.05; ** = *p* < 0.01; and *** = *p* < 0.001. Significance was assessed by ANOVA followed by Holm-Šídák test for multiple comparisons.

Together, these data demonstrate that Pf4 is associated with diminished neutrophil influx in this model in conjunction with reduced production of pro-inflammatory cytokines.

### Pf4 downregulates a distinct set of neutrophil chemokines in a TLR3- and IFNAR-dependent manner

To obtain a more complete picture of Pf4-induced cytokine changes to the bacterial stimulation, we moved to an *in vitro* system with LPS as the source of inflammation. We examined supernatants from human U937 macrophages treated with LPS with or without Pf4 phage via multiplexed bead-based assay (Figure 2A). We used U937 cells for these experiments because of the central role of macrophages in airway inflammation (*Byrne* et al., 2015; *Hashimoto* et al., 1996).

**Figure 2:**
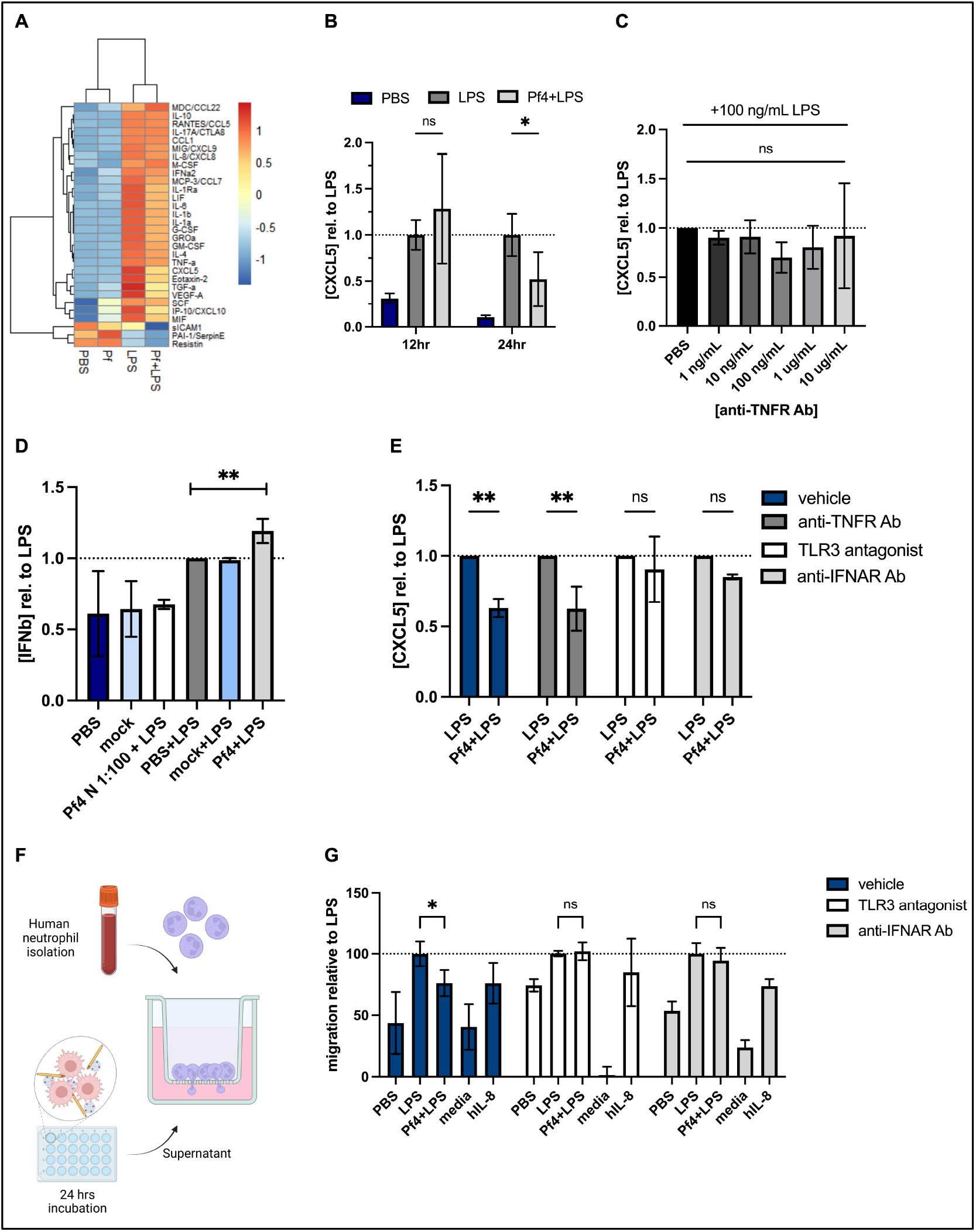
Pf4 impairs the production of a distinct set of chemokines in human macrophages which promote neutrophil migration in a TLR3- and IFNAR-dependent manner. (A) Luminex analysis of U937 macrophage supernatants treated with 100 ng/ml LPS and/or 10^9^ pfu/ml Pf4 for 24 hours. (B and C) CXCL5 protein levels of U937 macrophages incubated with 100 ng/ml LPS, or 100 ng/ml LPS + 10^9^ pfu/ml Pf4 at indicated time points. Anti-TNFR antibody was added in various concentrations to human U937 macrophages along with 100 ng/ml LPS and CXCL5 production was determined via ELISA. (D) U937 human macrophages were supplemented with PBS, mock Pf4 preparation, or 10^9^ pfu/ml Pf4 without or with 100 ng/ml LPS for 24 hours. IFNβ production was determined by ELISA. (E) U937 cells treated with vehicle (PBS), 100 ng/ml anti-TNFR antibody, 27 μM of a TLR3/dsRNA complex inhibitor, or 100 ng/ml anti-IFNAR antibody along with LPS in absence or presence of Pf4 for 24 hours were assessed for CXCL5 production by ELISA. (F) Schematic of human neutrophil migration assay. Human U937 macrophages were stimulated with LPS, Pf4 or TLR3 antagonists for 24 hours. The supernatant was isolated and used as a chemoattractant in a transwell setup. Human neutrophils isolated from freshly drawn blood and stained with Calcein AM, then applied to the top of a transwell plate with 3 μm pore size (STAR methods). Neutrophil migration towards conditioned media was assessed at 60 minutes. (G) Neutrophil migration normalized to LPS control. All data shown as the mean ± SD of at least 3 biological replicates with n ≥ 2 technical replicates per condition and experiment. * = *p* < 0.05; ** = *p* < 0.01; and *** = *p* < 0.001. Significance was assessed by ANOVA followed by Holm-Šídák test for multiple comparisons.

We observed that Pf4 downregulated production of multiple factors made in response to LPS including GROa/CXCL1, CXCL5, and TNF-α, as reported above, as well as other key pro-inflammatory proteins such as IL-6, IL-1a, IL-1b, and GM-CSF (Figure 2A). We chose to explore the functional impact of Pf4 on LPS-mediated production of CXCL5, a neutrophil chemoattractant, because dysregulation of neutrophil influx has been shown to strongly impact responses to *P. aeruginosa* and based on our findings in the acute *P. aeruginosa* lung model (*Koh* et al., 2009; *Lavoie* et al., 2011). Moreover, neutrophil influx is critical to control and clear *P. aeruginosa* infections (*Koh* et al., 2009; *Lavoie* et al., 2011).

We established that CXCL5 made in response to LPS is significantly downregulated by Pf4 at the protein level by 24 hours in U937 macrophages, analogous to the effect of Pf4 phage on TNF-α production reported previously (*Sweere* et al., 2019) (Figure 2B). We then sought to determine whether CXCL5 levels reflect impaired TNF-α signaling in this system. To that end, we blocked the TNF-α receptor (TNFR) prior to treatment with Pf4. We found that CXCL5 protein expression is not significantly attenuated by diminished TNF-α signaling (Figure 2C).

We previously reported that TNF-α production by macrophages was inhibited by Pf phage in a TLR3 and type 1 interferon-dependent manner (*Sweere* et al., 2019). We asked whether CXCL5 inhibition was also mediated in this way. We indeed observed that Pf4 induces type I IFN production by macrophages (Figure 2D). We then assessed the dependency of the present system on TLR3 by treating U937 macrophages with a small-molecule compound that specifically prevents dsRNAs from binding TLR3 (*Cheng* et al., 2011), and found that CXCL5 downregulation in response to Pf4 was abolished (Figure 2E). Similarly, cells treated with a blocking antibody against the type I IFN receptor, IFNAR1/2, showed a loss of phenotype (Figure 2E). Taken together, these data indicate that Pf4 downregulates multiple pro-inflammatory cytokines in human macrophages, including neutrophil chemoattractants, in a TLR3- and IFNAR-dependent manner. Moreover, the pathway that results in CXCL5 inhibition is parallel to but not dependent on the effects of Pf4 we previously reported on TNF-α (*Sweere* et al., 2019).

### Pf4 impairs neutrophil recruitment by human macrophages

To test the functional consequences of reduced chemokine production, we adapted a neutrophil migration assay (*Frevert* et al., 1998) using freshly isolated human primary neutrophils from multiple donors and conditioned media from human U937 macrophages. Macrophages were exposed to purified Pf4 preparation along with LPS for 24 hours to induce inflammatory cytokine production, and conditioned media from these cells was used as a chemoattractant in a transwell setup (Figure 2F). Neutrophils were stained with the live-cell dye calcein AM and tracked as they migrated through the 3 μm pores of the transwell insert.

We found that conditioned media from cells treated with a combination of Pf4 and LPS induced less migration than cells treated with LPS alone (Figure 2G), indicating that the presence of purified Pf4 is sufficient to alter macrophage function in response to bacterial stimulation. Inhibition of either TLR3 or IFNAR in macrophages resulted in loss of Pf4-driven reduced neutrophil chemoattraction as compared to the LPS control (Figure 2G).

Together, these data indicate that Pf4 reduces the ability of LPS to stimulate macrophages to induce granulocyte migration.

## DISCUSSION

We report that Pf4 phage downregulates multiple LPS-induced factors, including the potent neutrophil chemoattractant CXCL5. These suppressed macrophages are less effective at inducing neutrophil migration in a mouse model of acute *P. aeruginosa* lung infection, as well as *in vitro* using a human neutrophil migration assay. Neutrophil influx to sites of *P. aeruginosa* infection is critical to control and clear an infection (*Koh* et al., 2009; *Lavoie* et al., 2011). The findings we present in this work indicate that the presence of Pf4 phage could be a marker of negative outcome in patients infected with *P. aeruginosa*, through ineffective macrophage activation and subsequent impaired recruitment of neutrophils at early stages of infection. The intriguing question of whether Pf4-associated negative outcomes are related to the titres of Pf4 in the CF airways would be important to investigate further.

Although we chose to focus on the interaction between macrophages and neutrophils due to the observed downregulation of several neutrophil chemoattractants, several other cytokines altered by Pf4 stimulation may affect the course of an immune response to *P. aeruginosa*. In particular, the IL-1α/β and GM-CSF axis has been shown to be important for neutrophil longevity and thereby effective bacterial clearance (*Bober* et al., 1995; *Fossati* et al., 1998; *Laan* et al., 2003). In addition, GM-CSF has been shown to amplify bystander phagocyte cytokine production in *Legionella* infection (*Liu* et al., 2020).

The co-existence of filamentous phages with the bacterial hosts they infect suggests a potential symbiotic relationship. Indeed, prior work has demonstrated that Pf4 impacts *P. aeruginosa* pathogenesis during chronic pulmonary and wound infections (*Secor* et al., 2015; *Secor* et al., 2017; *Burgener* et al., 2019; *Sweere* et al., 2019). Unlike lytic phages which lyse the bacterial host cell, chronically infecting phages like Pf4 integrate their genetic material into the bacterial genome and co-exist with the bacteria in high titres. This co-existence assures that continued interaction between the phage and the bacterial cell will continue, manifesting in such phenomena as liquid crystal formation using bacterial polymers and phage-formed occlusive sheaths that protect bacterial cells from antibiotics (*Secor* et al., 2015; *Tarafder* et al., 2020). In this work, we present yet another instance of *Pf4-Pseduomonas* symbiosis, in which the phage prevents the clearance of the bacterium by reducing phagocyte immune responses.

## ACKNOWLEDGMENTS

This work was funded by the grants from the NIH including R01 HL148184-01, R01 AI12492093, and R01 DC019965, as well as grants from the Cystic Fibrosis Foundation and the Emerson Collective. Funding support was also received from the National Science Foundation Graduate Research Fellowship and the Gabilan Stanford Graduate Fellowship. Graphical schematics created with BioRender.com.

## Notes

### Competing Interest Statement

The authors have declared no competing interest.

